# AbLang: An antibody language model for completing antibody sequences

**DOI:** 10.1101/2022.01.20.477061

**Authors:** Tobias H. Olsen, Iain H. Moal, Charlotte M. Deane

**Affiliations:** Department of Statistics, University of Oxford, Oxford OX1 3LB, United Kingdom; GSK Medicines Research Centre, GSK, Stevenage SG1 2NY, United Kingdom

## Abstract

**Motivation:** General protein language models have been shown to summarise the semantics of protein sequences into representations that are useful for state-of-the-art predictive methods. However, for antibody specific problems, such as restoring residues lost due to sequencing errors, a model trained solely on antibodies may be more powerful. Antibodies are one of the few protein types where the volume of sequence data needed for such language models is available, for example in the Observed Antibody Space (OAS) database.

**Results:** Here, we introduce AbLang, a language model trained on the antibody sequences in the OAS database. We demonstrate the power of AbLang by using it to restore missing residues in antibody sequence data, a key issue with B-cell receptor repertoire sequencing, over 40% of OAS sequences are missing the first 15 amino acids. AbLang restores the missing residues of antibody sequences better than using IMGT germlines or the general protein language model ESM-1b. Further, AbLang does not require knowledge of the germline of the antibody and is seven times faster than ESM-1b.

**Availability and Implementation:** AbLang is a python package available at https://github.com/oxpig/AbLang.

## 1 Introduction

Recent progress within protein informatics has led to the development of pre-trained protein representations, derived from protein language models such as ESM-1b (Rives *et al*., 2021), which have been used to perform state-of-the-art predictive tasks. Such protein language models require vast amounts of training data and so far have tended to use all protein sequences and therefore be general protein representations (Alley *et al*., 2019, Elnaggar *et al*., 2021, Rives *et al*., 2021). With the creation of the Observed Antibody Space (OAS) database (Kovaltsuk *et al*., 2018) and subsequent update (Olsen *et al*., 2021), enough curated antibody sequences are now available to train a language model specifically for antibodies. An antibody specific model that learnt the semantics of their sequences would allow for more precise predictions of antibody properties and new use cases.

Over the last decade, billions of antibodies have been sequenced (Chaudhary and Wesemann, 2018). However, in some cases the sequenced antibodies are missing residues due either to sequencing errors, such as ambiguous bases (Huse *et al*., 2007), or the limitations of the sequencing techniques used (Kim and Park, 2019). We find that in OAS, ~80% of the sequences are missing more than one residue at the N-terminal and ~43% the first 15 positions, and ~1% contain at least one ambiguous residue. The ability to accurately restore these missing residues would increase data availability and be of benefit to antibody drug discovery. Currently, sequence imputation can only be done by identifying the correct ImMunoGeneTics (IMGT) germlines from the IMGT/GENE-DB (Giudicelli *et al*., 2005) and using the germline sequence to add the missing residues. This approach requires correctly determining the allele of the sequence, a process that can be time consuming and/or produce ambiguous results.

Here, we present AbLang, an antibody specific language model trained on either the heavy or light chain antibody sequences from OAS. While AbLang can be used to create representations for residue or sequence specific predictions and residue engineering, in this paper we focus on showing how AbLang can be used to accurately restore missing residues in antibody sequences, more accurately than using IMGT germlines or a general protein model like ESM-1b.

## 2 Materials and methods

Two AbLang models were trained, one for heavy and one for light chains. Each AbLang model consists of two parts: AbRep, which creates representations from antibody sequences, and AbHead, which uses the representations to predict the likelihood of each amino acid at each position (Figure 1).

**Figure 1:**
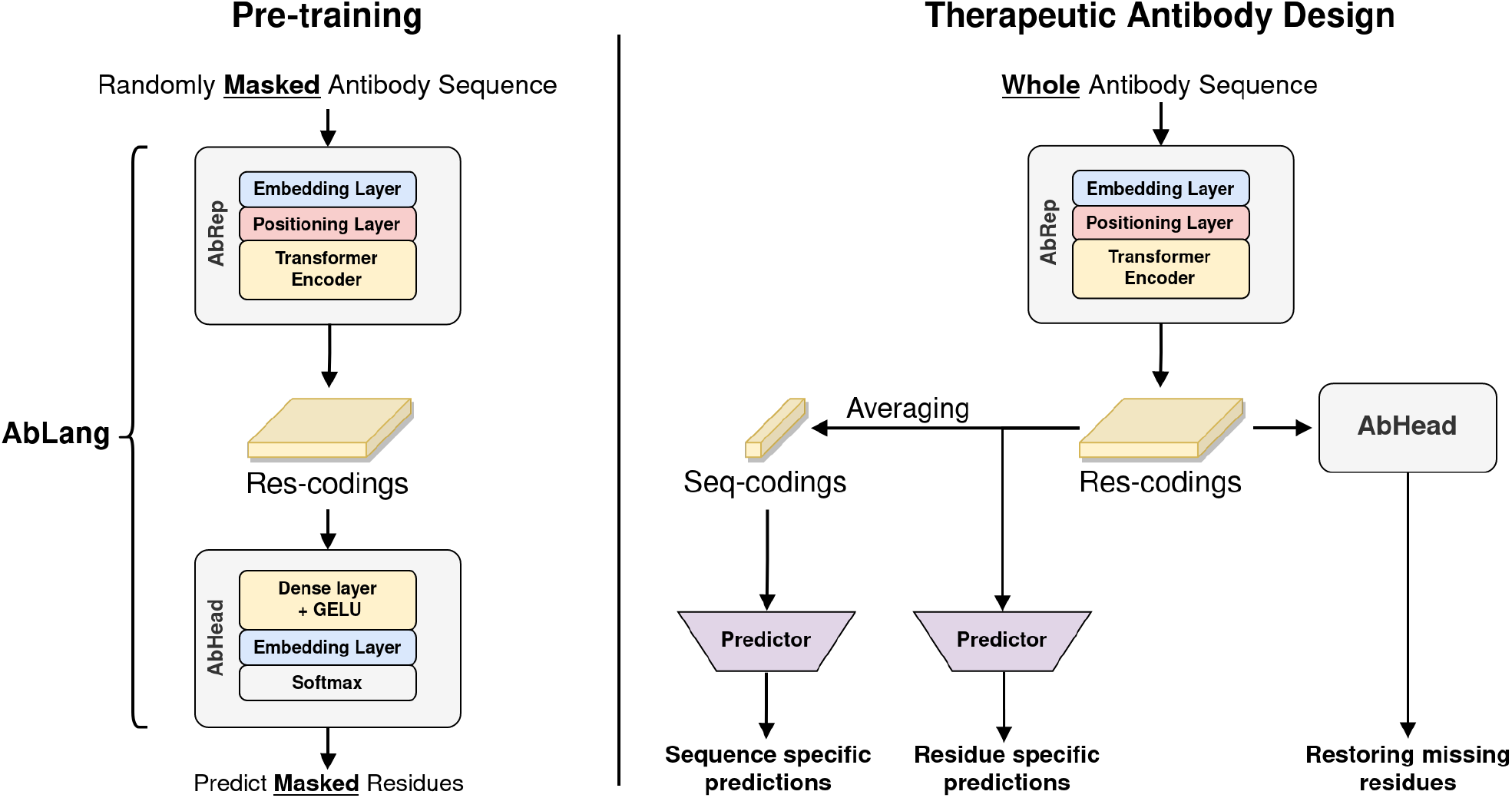
Overview of the architecture of AbLang, AbRep and AbHead, and examples of possible use cases. For pre-training, residues are randomly masked in each sequence and the masked residues are predicted and compared to the original residue. After pre-training, the model can, through different use cases, be used to improve therapeutic antibody design. AbHead can be removed and the res-codings from AbRep used for either residue or sequence specific predictions, or AbHead can be kept and used for restoring missing residues or exploring possible mutations.

AbLang was implemented using PyTorch 1.8.1 and was inspired by HuggingFace’s (Wolf *et al*., 2020) Transformer 3.0.2 library. AbRep follows the architecture of RoBERTa (Liu *et al*., 2019), except it uses a learned positional embedding layer with a max length of 160. Each of its 12 transformer blocks has 12 attenuated heads, an inner hidden size of 3072 and a hidden size of 768. From AbRep, the res-codings (768 values for each residue) are obtained. AbHead follows the design of RoBERTa’s head model, with a hidden size of 768.

During training, between 1-25% of residues from each sequence were selected, and of these, 80% were masked, 10% randomly changed to another residue and 10% left unchanged. One AbLang model was trained on heavy chain sequences for 20 epochs with a batch size of 8,192, and another on light chain sequences for 40 epochs with a batch size of 4,096. Both models were optimized using an Adam optimizer with a linear warm-up period for 5% of the steps, a peak learning rate of 0.0002, a learning rate decrease following a cosine function, and a weight decay of 0.01. For every dropout and layer normalization, a 0.1 rate and 1e^−12^ epsilon was used. The hyperparameters were selected to be similar to those used in the RoBERTa paper (Liu *et al*., 2019).

## 3 Results

### 3.1 Data preparation

All antibody sequences seen three or more times in the OAS database as of Oct. 2021 were downloaded. The heavy and light chain sequences were then clustered separately based on identical CDR3s and thereafter clustered further by 70% identity over the whole sequence using Linclust (Steinegger and Soding, 2018), with the longest sequence selected from each cluster. The selected sequences were then randomly divided into training sets of 14,126,724 heavy and 187,068 light sequences, and two evaluation sets of 100,000 heavy and 50,000 light sequences. The training sets were then used to train AbLang as described in the Materials and methods section.

### 3.2 AbLang’s antibody sequence representations

AbLang can be used to generate three different representations of antibody sequences. The first representation, the res-codings, consists of 768 values for each residue, useful for residue specific predictions. The second representation, the seq-codings, represent the whole sequence and is derived from the mean of all res-codings in a sequence. The seq-codings are 768 values for each sequence and are useful for sequence specific predictions. Additionally, they have the benefit of having the same length for each sequence, removing the need to align antibody sequences. Lastly, AbLang can be used to generate the likelihoods of each amino acid at each position in a given antibody sequence, useful for antibody engineering.

To investigate the sequence information extracted by AbLang and compare it to that of ESM-1b, we visualised the AbLang and ESM-1b sequence representations of 10,000 naïve and 10,000 memory B-cell sequences from Ghraichy et al (2021) with a t-SNE (van der Maaten and Hinton, 2008) plot (Figure 2).

**Figure 2:**
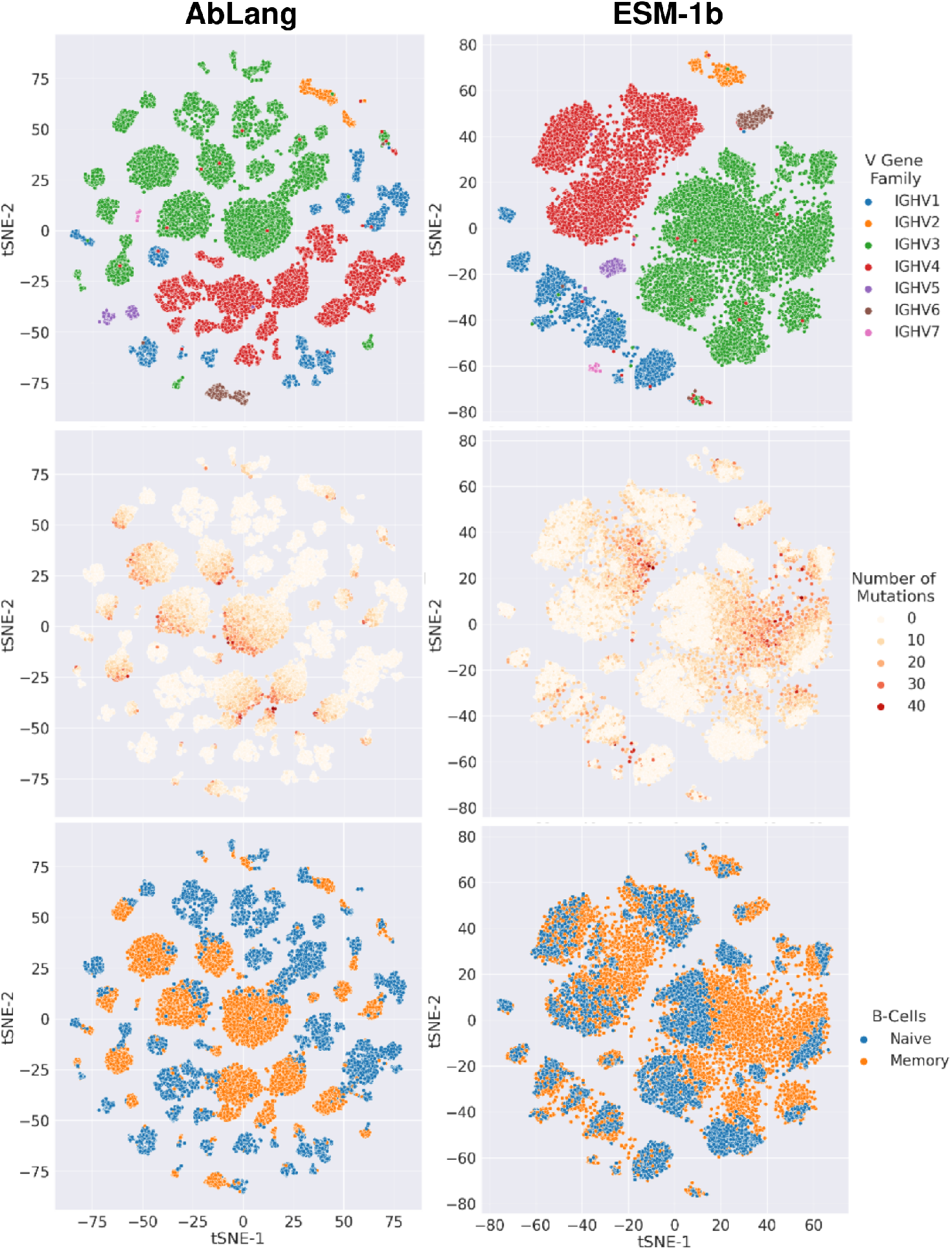
Comparison of AbLang and ESM-1b representations at clustering sequences based on their V-genes, originating cell type and number of mutations.

As seen in Figure 2, AbLang and ESM-1b both separate the sequences based on their V-genes, however, AbLang separates the V-genes into smaller clusters. Within the AbLang clusters, a clearer separation can be seen between naive B-cells and memory B-cells than with ESM-1b’s clusters. Further, the memory B-cells, in AbLang’s clusters, appear to be ordered based on a gradual increase in mutations. This could indicate that the AbLang representations contain information about the order of antibody mutations.

### 3.3 AbLang for restoring missing residues

AbLang’s representations can be used for a plethora of antibody design applications. As an example, we use AbLang to restore missing residues in antibody sequences. Figure 3 demonstrates the need for such a tool, showing how over 40% of the sequences in OAS are missing the first 15 residues and ~80% of the sequences are missing more than one residue at the N-terminal.

**Figure 3:**
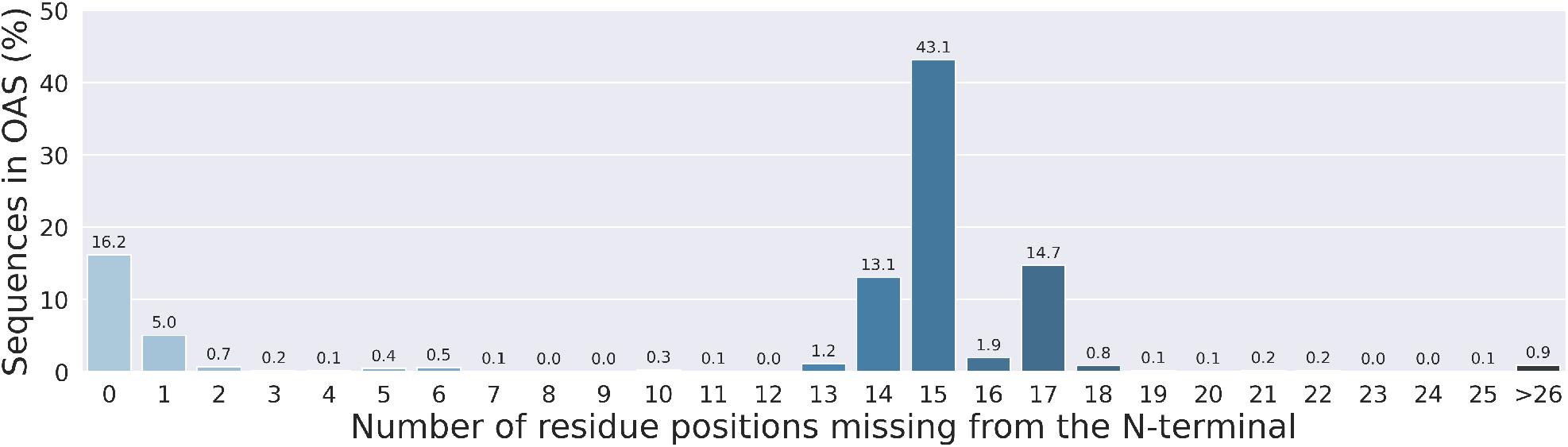
Overview of the antibody sequences in OAS, showing the percentage of sequences and number of residues they are missing from the N-terminus. Over 40% of the sequences in OAS are missing the first 15 residues.

The input to AbLang for sequence restoration is an antibody sequence with asterisks for unknown residues (Figure 4). AbLang restore the missing residues by predicting the likelihood of each amino acid at the marked positions, with the amino acid with the highest likelihood then selected as the prediction.

**Figure 4:**
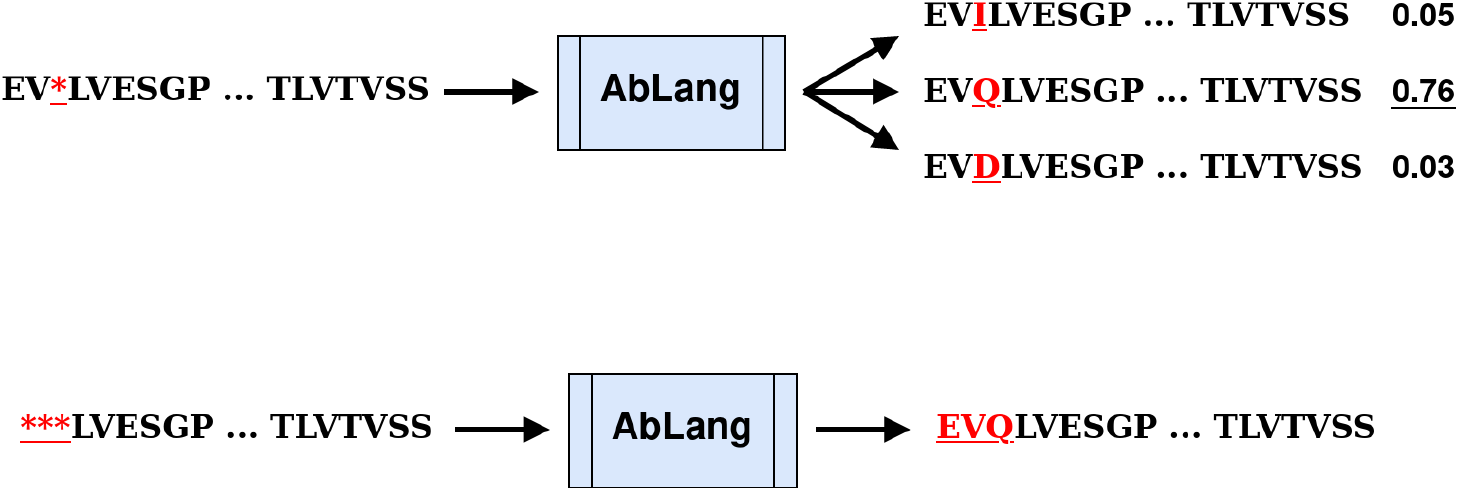
Illustration of how AbLang restores missing residues. An input sequence has asterisks at the positions to predict. AbLang predicts the amino acid with the highest likelihood.

We tested the ability of AbLang to restore both the N-terminal of antibody sequences and missing residues randomly scattered throughout the sequence. From the evaluation sets, 100 complete sequences for each of the 20 heavy and 42 light human V alleles seen in the evaluation set were randomly selected. These 2000 heavy and 4200 light sequences were used as the test set.

Figure 5 shows a comparison of AbLang, to the general protein model ESM-1b and to the use of germline residues for the prediction of missing residues in an antibody sequence. Sequences were numbered in the IMGT scheme using ANARCI (Dunbar and Deane, 2016) and positions from 1 up to 30 were masked and then restored using the three different methods. The accuracy of this restoration was measured as the percentage of correctly predicted amino acids. IMGT germlines and AbLang achieve comparable accuracy, both restore missing N-terminal residues with accuracies of around 96% and 98% for the first 15 positions of the light and heavy chain, respectively. ESM-1b achieves far poorer performance achieving accuracies of 54% and 64%. The performance of IMGT germlines and AbLang are very similar, but the IMGT germline method requires knowledge of or accurate prediction of the germline, while AbLang can be rapidly deployed without any additional information.

In some cases, sequencing errors can result in residues being unknown at random sites throughout the antibody sequence. The ability of AbLang, IMGT germlines and ESM-1b to predict residues at randomly selected positions was also compared. Using the same test set as above, one, five or ten residues were randomly masked in each sequence’s V-region. AbLang is more accurate at this task than both IMGT germlines and ESM-1b for both heavy and light chains. AbLang is also the fastest of the three methods, able to process 100 sequences in 6.5 seconds to ESM-1b’s 44.9 seconds, using 4 cores on an Intel Core i7-10700.

**Figure 5:**
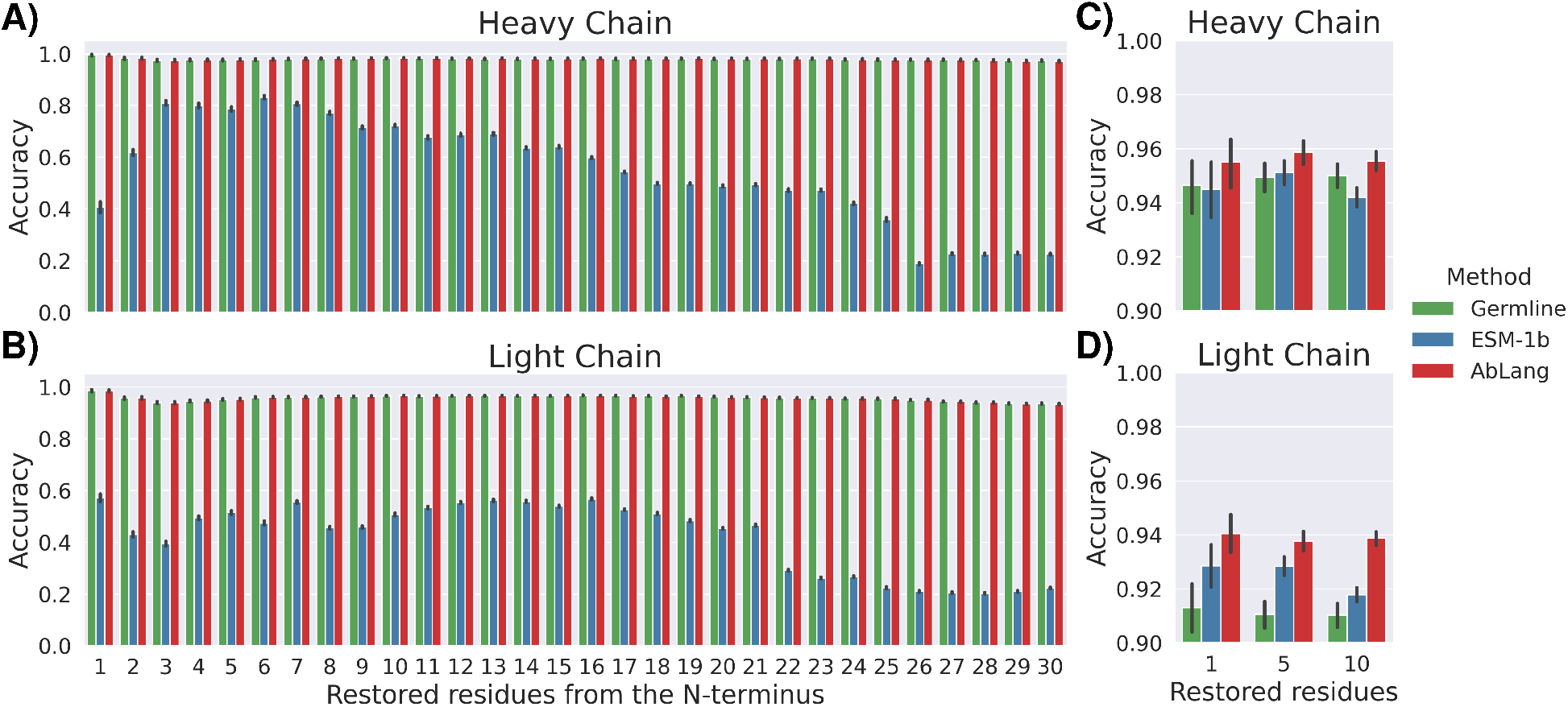
Antibody sequence restoration using IMGT germline sequences (green), a general protein language model ESM-1b (blue), and the antibody specific language model AbLang (red). A-B, shows the restoration of sequences missing up to 30 residues of the N-terminal, and C-D, the restoration of sequences with a random set (one, five or ten) of missing residues.

## 4 Discussion

A language model specific for antibodies, should learn a deeper understanding of the semantics of antibodies than current general protein language models. In this work we present AbLang, a language model trained on a large dataset of antibody sequences from OAS. To demonstrate that AbLang has learnt a useful representation of antibodies, we show how AbLang’s sequence representations contain knowledge of the germlines, originating cell type and number of mutations.

We also demonstrate the use of AbLang to restore missing residues in antibody sequences, and show how AbLang performs on par or better than using IMGT germlines, but without the need to have knowledge of the germline. Further, we show how AbLang restores residues more accurately and faster than a current state-of-the-art protein language model ESM-1b, emphasising the benefits and potential of an antibody specific language model.

